# Deep-learning-based flexible pipeline for segmenting and tracking cells in 3D image time series for whole brain imaging

**DOI:** 10.1101/385567

**Authors:** Chentao Wen, Takuya Miura, Yukako Fujie, Takayuki Teramoto, Takeshi Ishihara, Koutarou D. Kimura

**Author notes:** Correspondence:Chentao Wen.

## Abstract

The brain is a complex system that operates based on coordinated neuronal activities. Brain-wide cellular calcium imaging techniques have quickly advanced in recent years and become powerful tools for understanding the neuronal activities of small animal models. The whole brain imaging generally requires to extract the neuronal activities from three-dimensional (3D) image series. Unfortunately, the 3D image series are obtained under imaging conditions different among laboratories and extracting neuronal activities from the data requires multiple processes. Therefore researchers need to develop their own software, which has prevented the application of whole-brain imaging experiments in more laboratories. Here, we combined traditional image processing techniques with the powerful deep-learning method which can be flexibly modified to fit 3D image data in the nematode *Caenorhabditis elegans* obtained under different conditions. We first trained the 3D U-net deep network to classify each pixel into cell and non-cell categories. Cells merged as a whole region were further separated into individual cells by watershed segmentation. The cells were then tracked in 3D space over time with the combination of a feedforward network and a point set registration method to use local and global relative positions of the cells, respectively. Remarkably, one manually annotated 3D image combined with data augmentation was sufficient for training the deep networks to obtain satisfactory tracking results. Our method correctly tracked more than 98% of neurons in three different image datasets and successfully extracted brain-wide neuronal activities. Our method worked well even when the sampling rate was reduced: 86% correct in case 4/5 frames were removed, and when artificial noise was added into the raw images: 91% correct in case 35 times of background-level noise was added. Our results proved that deep learning is widely applicable to different datasets and can help us in establishing a flexible pipeline for extracting whole brain activities.

## 1. Introduction

The brain is a complex system consisting of thousands of millions of neurons that interact with each other in a highly organized way [1–4]. To understand how this complex system works, we need to monitor whole-brain neuronal activity on living animals [5,6]. Traditional electrophysiological techniques such as multichannel extracellular recording can only measure the simultaneous activities of a small proportion of neurons. While these types of studies can be used to elucidate local features of specific neurons, they cannot help us understand the brain as a whole. Other techniques such as electroencephalography (EEG), magnetoencephalography (MEG), and functional magnetic resonance imaging (fMRI) can monitor whole brain activities. However, those recorded signals reflect activities from a mass of neurons but not individual neuronal activity [7, 8]. On the other hand, recent whole-brain calcium imaging techniques have achieved cellular level resolution. These techniques have been used in small animals with transparent brains, such as larval zebrafish [9–13] and the nematode *Caenorhabditis elegans* [14–18].

Although multiple studies have been reported, extracting neuronal activities from whole-brain images is still challenging. Because a whole-brain image time-series could contain a large number of neurons across hundreds or thousands of frames, manually marking and tracking the neurons would be impossible. Thus, automatic methods for processing such 3D images are required. In previous studies [15,16,18,19], however, different labs have used different strains, different microscopy systems, different imaging parameters, and different constraint conditions; consequently, each lab needed to develop different methods to process their whole-brain 3D images, which is time-consuming and labor intensive. Thus, a flexible method that can be applied for different imaging conditions is highly desired. There are several benefits of using a flexible method: 1. Easy modification will allow researchers to quickly start a whole-brain imaging experiment; 2. Imaging conditions can be freely changed without requiring scientists to develop a new method; 3. Datasets obtained from different labs can be uniformly analyzed; and 4. Researchers do not need to learn multiple methods for image processing.

Traditionally, image processing algorithms are designed for appropriate features of the image to solve a specific task [20]. However, such manually designed methods cannot be easily applied to images generated in very different conditions. In recent years, deep-learning techniques have become popular in image processing tasks. Deep-learning methods have out-performed traditional methods in some image processing tasks such as image classification [21], thus demonstrating the power of this technique. Moreover, deep-learning methods can be flexibly modified to perform with very different image conditions.

Deep-learning methods use an artificial neural network with multiple layers, i.e., a deep network, to process data such as images. Each basic unit in the deep network computes the weighted sum of its inputs from previous layer and passes the result to a non-linear function [22]. By connecting such basic units into a large network, we can approximate a variety of complex functions [23], such as transforming an image to a different style. Deeper networks with more layers can recognize more complex/global features (e.g., person, car, cell) by combining simple/local features (e.g., corners, contours) [24]; thus, deeper networks can solve difficult image processing tasks. Even though these complex features are rather difficult to design using traditional methods, deep networks can learn these features automatically from data. Despite these advantages, deep-learning methods have not been used for processing 3D whole-brain images in previous studies.

In this study, we developed a flexible method for extracting neuronal activities from whole-brain images of *C. elegans*. In the proposed method, powerful and flexible deep-learning techniques are combined with traditional image processing techniques to process 3D whole-brain image time-series. The deep-learning parameters can be automatically learned, and the traditional techniques in our method have significantly fewer parameters compared to previously reported methods [16] (Table 1). In this way, our proposed method can be more easily modified to fit different imaging conditions. We used two specific deep networks: the 3D U-net [25], and a fully connected feedforward network. Both networks can be efficiently trained using only one manually annotated 3D image frame, which reduces annotation time. Using our new method, we successfully extracted activities from three types of datasets, which were obtained under different imaging conditions. Our method also successfully extracted neuronal activities from more difficult image time-series that were generated by adding noise or by removing intermediate frames from a dataset. Our results demonstrated that deep learning could be used to establish a flexible method for extracting neuronal activities; thus, such algorithms could be used by more laboratories performing whole-brain calcium imaging without considerable modification.

## 2. Results

### 2.1 Objective for the proposed algorithm

Our primary objective was to extract neuronal activities from whole brain 3D images. A *C. elegans* brain (Fig. 1A) was scanned under the microscope from the lowest layer to the highest layer (Fig. 1B), and the head neurons were marked by a fluorescent protein. Each set of the scanned 2D images from bottom to top formed one frame of the 3D image. Multiple cycles of scanning formed the 3D image time-series (Fig. 1B; 1C top). The size of a typical 3D image (see type 1 dataset) was around 80 μm in width, 170 μm in length, and 40 μm in depth, covering the entire head region of a young adult worm. The time interval between two successive 3D images was 1 s or less, and the time series usually lasted for several hundreds of seconds. Two types of 3D images were simultaneously scanned: 1. the neuron nucleus markers, and 2 the calcium indicators, which measured the concentration of the calcium in each neuron, thus reflecting neuronal activities. To extract the neuronal activities at each frame, we needed to solve two major problems: 1. Each neuron must be assigned a specific label, and 2. Each neuron must be tracked over time. The process of assigning a label to each neuron is called segmentation (Fig. 1C, bottom left). Note that a single neuron can span different layers or different neurons can merge, forming a connected region, because of their close proximity. These factors can complicate the segmentation process. Moreover, due to the slight movement of the worms under the anesthetized/constraint condition, the position of each neuron changed during the imaging, and the positions of those neurons must be updated in a process known as tracking (Fig. 1C, bottom right). Both the segmentation and tracking processes were performed using the nucleus marker images. Then, the neuronal activities could be easily extracted by calculating the mean intensity of the calcium indicator at the regions corresponding to each neuron. To avoid movement artifacts, the extracted calcium signals were usually normalized, e.g., divided by the mean intensity of the nucleus marker.

**Figure 1.**
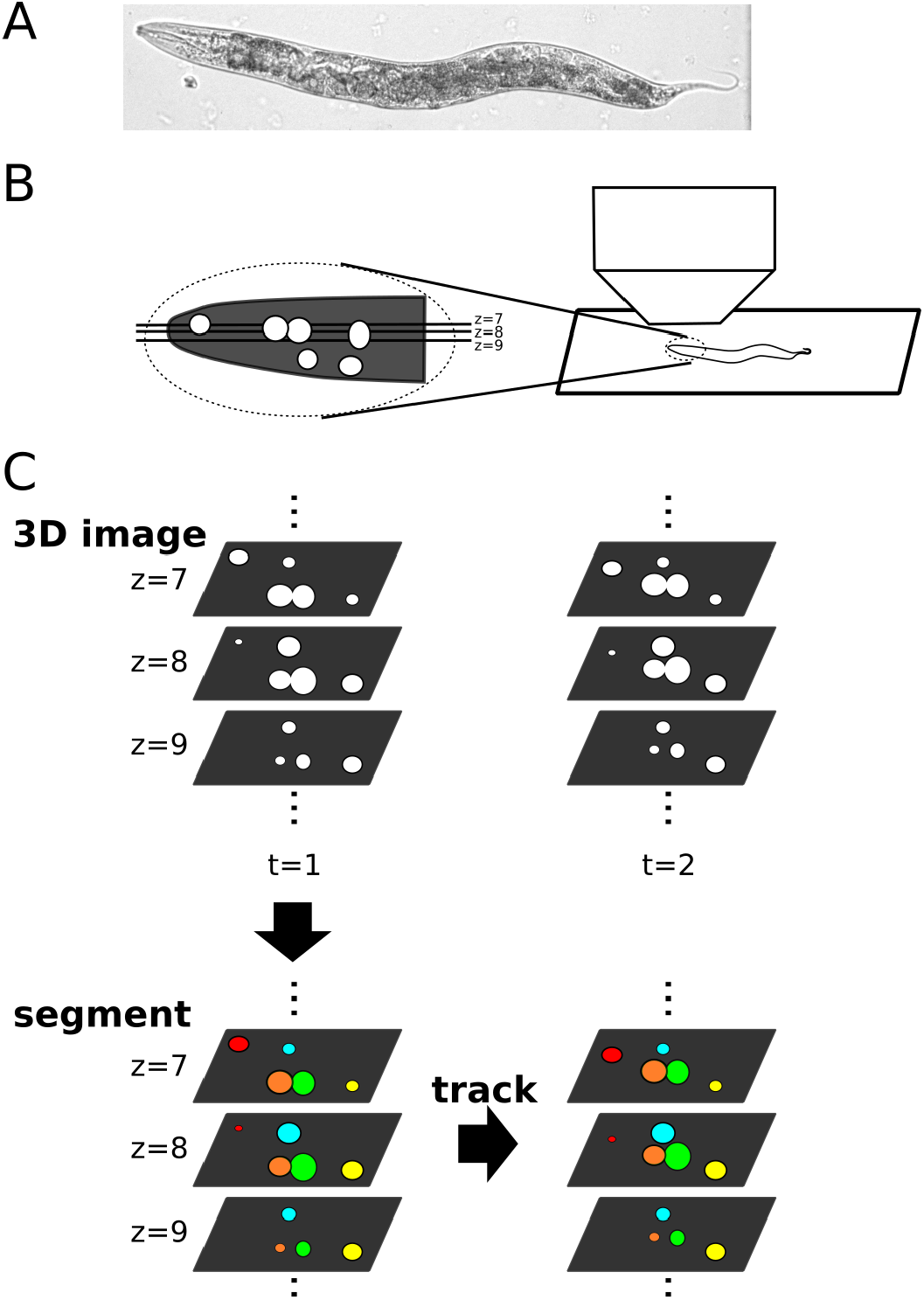
Extraction of neuronal activity from 3D data sets. A) Photograph of *C. elegans* taken using our microscopy system at a low amplification. B) Right: Schematic of the 3D imaging experiment. The plane parallel with the slide glass is the x-y plane, whose orthogonal direction is the z direction. Left: an illustration of head neurons (white circles) projected onto the x-z plane. C) Top: An schematic of 3D images received from the microscope. Each 3D image consists of a series of 2D images at different z levels. The microscope takes 3D images at different times and forms a 3D image time series. Bottom: Example of segmentation and tracking. We segmented the cell-like regions of a 3D image into individual cells, and then those cells are tracked over time to update their positions due to movements of the worm.

### 2.2 Overall procedure for the proposed method

Fig. 2 illustrates the overall procedure for the proposed method. After pre-processing (see Methods), we performed automatic segmentation in all 3D images using the 3D U-net + watershed (see texts below), and we only manually corrected mistakes in the first frame of the resulting segmented image. The manually confirmed cells were tracked, i.e., their positions were updated successively in following frames. To update their positions, we inferred a transformation function from each previous frame to its subsequent frame using a feedforward network + point set registration + accurate correction (see texts below).

**Figure 2.**
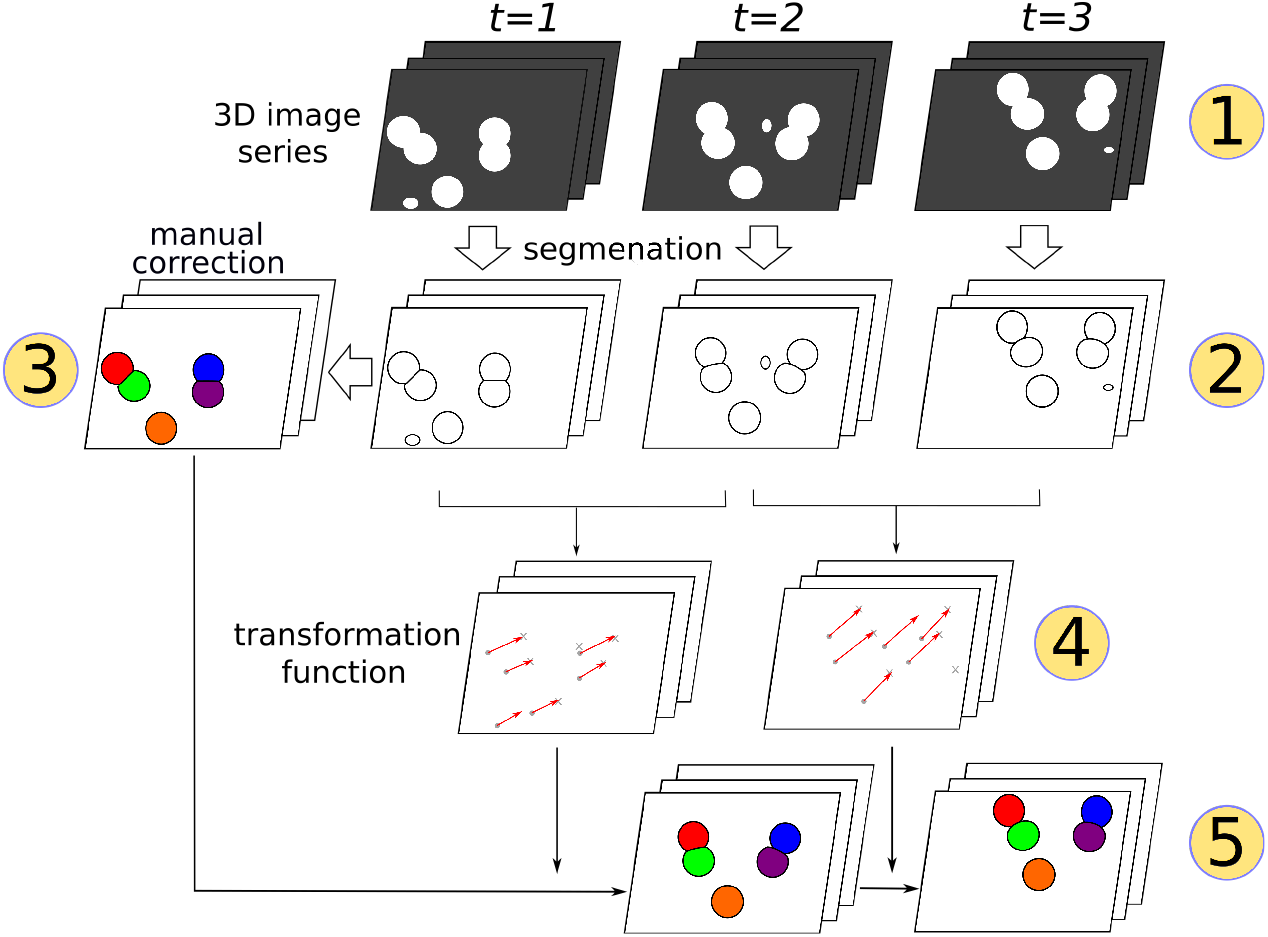
Overall procedure of our segmentation and tracking method. Pre-processed 3D images were automatically segmented into discrete regions. The first frame of the segmentation was manually corrected. In following frames, the positions of the manually confirmed cells were successively updated by inferred transformation functions. The circled numbers indicate the five different procedure steps: 1. Pre-processed 3D images; 2. Automatic segmentation; 3. Manual correction of the segmentation in the first frame; 4. Inferred transformation functions; and 5. Tracking results.

### 2.3 Segmentation

#### 2.3.1 Segmentation procedure

The raw image consisted of neurons with different intensities, sizes, shapes, and textures. Using the 3D U-net, we obtained cell-like regions that were close to those identified visually. Some cell-like regions included multiple neurons, so they were further separated into individual neurons using the watershed method [26]. We manually corrected the segmentation for the first frame of the image.

#### 2.3.2 3D U-net

Cell segmentation in 2D images using deep networks has been previously reported [27,28]. Instead of the more traditional convolutional neural network, which predicts only one pixel in one implementation of the network [27], we utilized a new structure called U-net, which can more-efficiently generate cell-like regions in the entire image using a single implementation of the network [28]. In real experiments, image series usually consist of hundreds or even thousands of images; thus, efficiency is essential. Both 2D and 3D versions of U-net have been proposed; however, only the 2D version has been previously tested in cell segmentation [25,28]. In this study, we utilized the 3D version of U-net for cell segmentation [25]. We modified the size of the input, output, and individual layers to suit images with different resolutions (Fig 3A, Fig. S1). The modification was performed because, depending on the resolution of the 3D image, each unit of the same U-net will have a different receptive fields. Using these modified structures, our method appropriately detected cell-like regions in all three test datasets as well as the generated datasets (see below).

**Figure 3.**
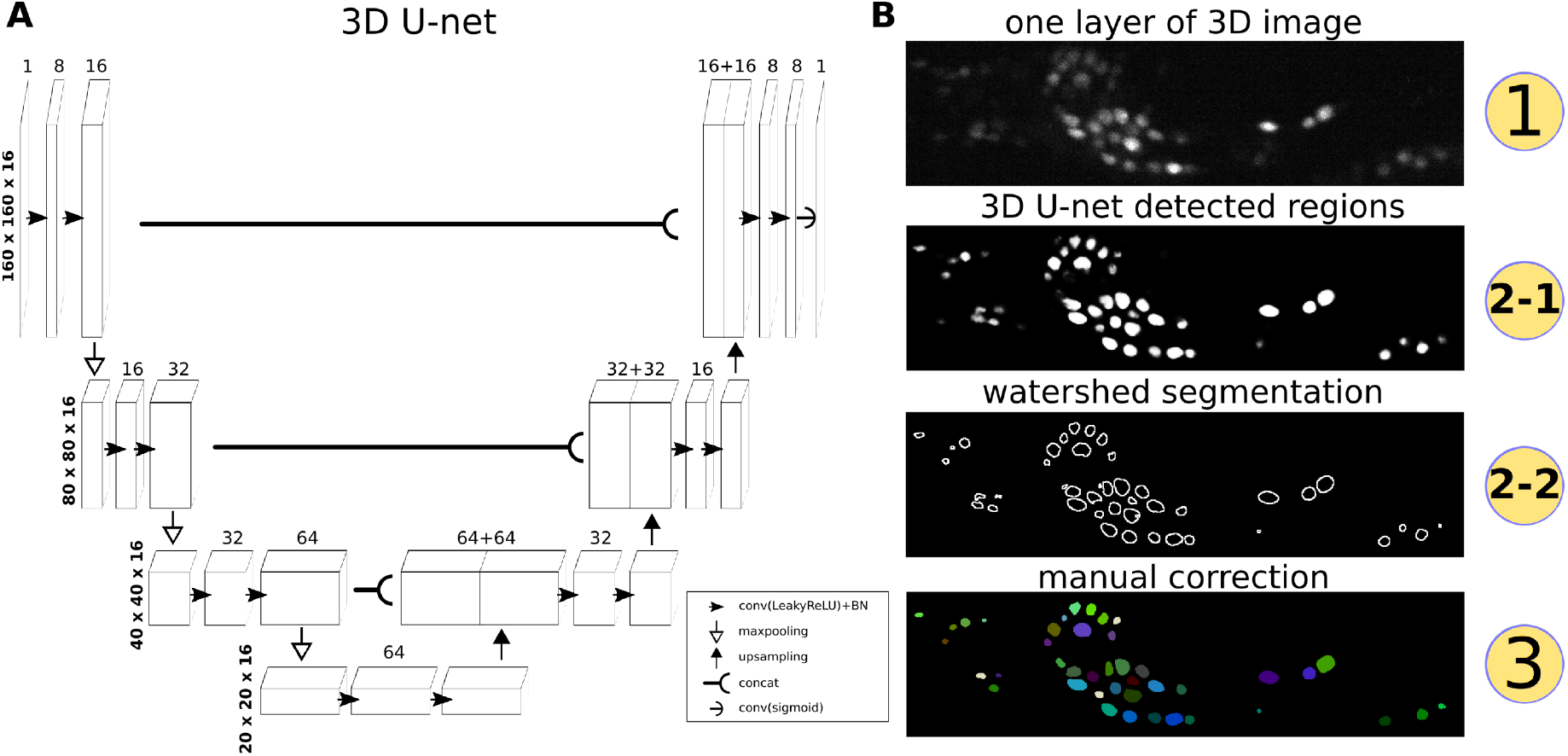
Procedure for segmentation with 3D U-net and post-processing. A) Structure of the 3D U-net. Numbers on each intermediate layer indicate the number of convolutional filters, while numbers at left of each row indicate the size of the 3D input, output, and intermediate layers. B) Schematic of the segmentation steps using one layer of a 3D image. The circled numbers correspond to the numbers in Figure 2.

The cell-like regions detected using the 3D U-net required a secondary step to separate the connected cells with the watershed method (see Method). The primary merit of using the 3D U-net is its transformation of different qualities of 3D images into images of the same form, which distinguished cell-like regions and background regions, thus making the proposed algorithm more robust to various imaging conditions. In next section, we explain how these segmented neurons in the 3D image time series are tracked over time.

### 2.4 Tracking

#### 2.4.1 Tracking procedure

There are multiple possible strategies for tracking neurons. One method is to utilize intensity or textural information of each neuron. Based on such information, we could update position of each neuron by searching for its new position one by one in an adjacent region in next frame. However, the problem with this strategy is also apparent: neurons cannot be tracked if their movements are greater than a certain threshold.

Another strategy is to use point set registration methods [29–31]. In these methods, the intensity and textural information of the neurons are ignored, and the neurons are represented only by their center points. The center points of all neurons in one frame constitute a point set. Point set registration methods can be used to transform the positions of a point set from one frame to the next, even when large movements arise. The problem with this strategy is that the transformations are not always accurate because the intensity and textural information identifying each neuron are ignored.

In this study, we combined the abovementioned strategies to obtain more consistent results (Fig. 4A). We applied a recently reported point set registration method that considers both global and local structures (PR-GLS) [31] to obtain a rough transformation from frame t to frame t+1. The PR-GLS method requires an initial match as a starting point, which we obtained using our feedforward network (see text below). The transformation function generated by PR-GLS was then corrected by utilizing the intensity/texture information contained in the 3D U-net output.

**Figure 4.**
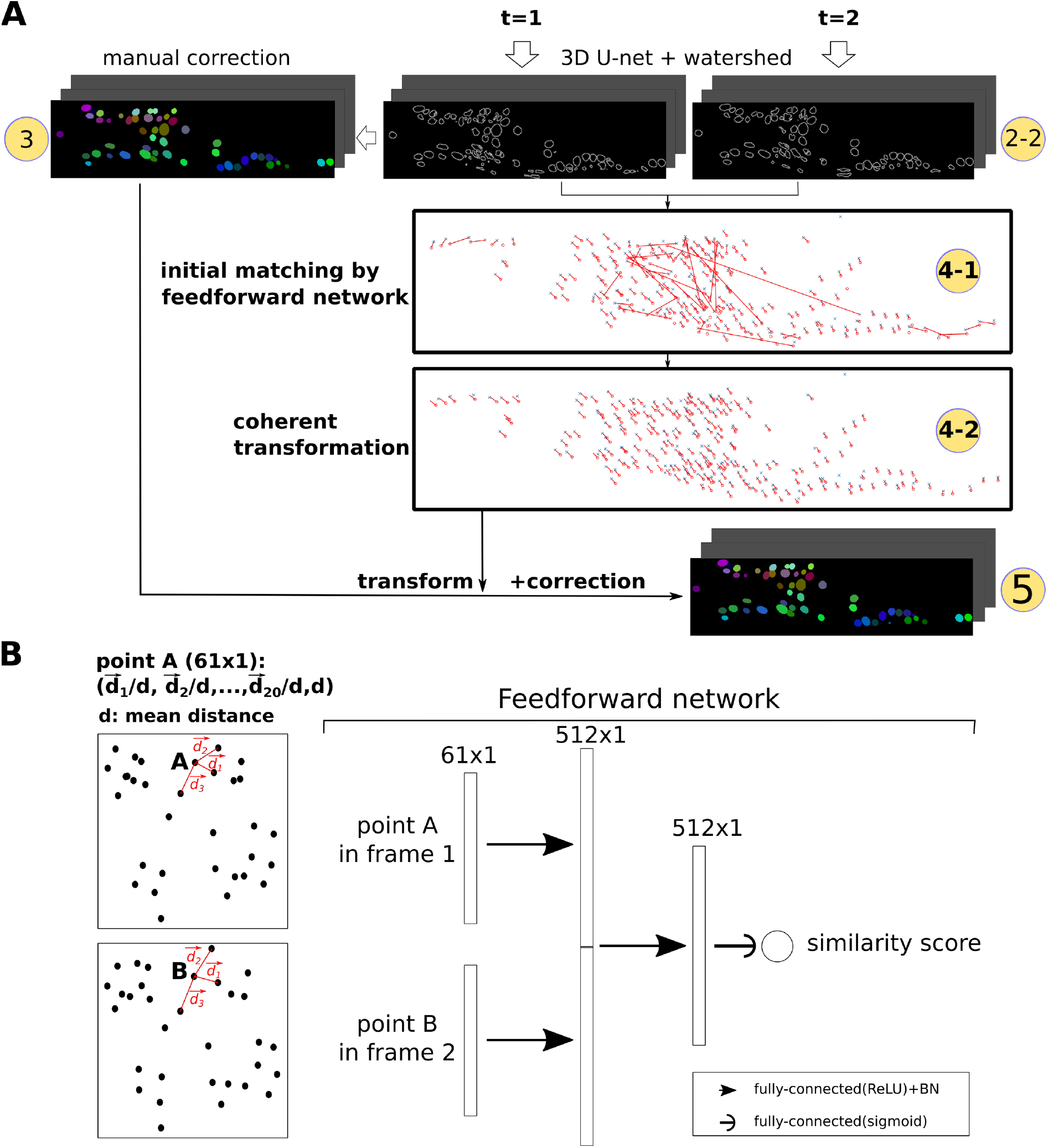
Tracking method. A) Procedures for the tracking method. The circled numbers correspond to those in Figure 2 and Figure 3. In 4-1 and 4-2, the red circles are the centers of segmented neurons at t=1; blue crosses are the centers at t=2; and the red lines are the inferred matching/transformation from t=1 to t=2. B) Left: Definition of the position of each point. Right: Structure of the feedforward network for calculating the similarity score between two points. Numbers on each layer indicate the shape of the layer.

#### 2.4.2 Initial matching performed by the feedforward network

The PR-GLS method requires an initial match, i.e., a set of correspondences between neurons in two frames. The correspondences can be estimated based on the positions of each neuron relative to other neurons, assuming that these relative positions do not significantly change even during the worm’s movement. By comparing the similarity of the relative positions of two neurons, we can determine whether they are the same neurons.

One traditional representation of relative positions is fast point feature histograms (FPFH) [32]. A previous study [31] successfully used the FPFH method for matching artificial point set datasets. However, we found that FPFH gave a poor initial match for the datasets considered in this study, perhaps because of the sparse distribution of the worm’s neurons (Fig. S2).

We thus designed a three-layer feedforward network to improve the initial match (Fig. 4B). The feedforward network is a basic but quintessential structure in deep-learning methods. In fact, both the convolutional neural network and U-net are specialized kinds of feedforward networks [24]. Our feedforward network successfully learned the appropriate representations of the relative neuron positions. By comparing the representations between two points, the network generated a similarity score between two neurons. We obtained better initial matching based on the similarity score than the matching by FPFH method (Fig. S2); however, a few incorrect matches were observed (Fig. 4A).

#### 2.4.3 Coherent non-rigid transformation by PR-GLS

The initial match obtained by the feedforward network usually had some incorrect matches (Fig. 4A), mainly because of problems in the automatic segmentation. In reality, the head of the worm cannot be deformed in an arbitrary way; therefore, nearby points should have coherent motions. By constraining the motions to be coherent, we can correct the initial match and obtain a more reliable transformation function. By applying the PR-GLS method [31] to the initial match, we obtained a more reliable transformation function in which all obvious incorrect matches were corrected (Fig. 4A).

#### 2.4.4 Accurate correction

The PR-GLS method usually generates coherent transformation functions very close to the real transformation. However, small differences still occur. Without correction, these small differences can cumulate over time to become large differences. Thus, we added one more correction step in which the center positions of each cell were moved slightly to the centers of each 3D U-net detected region (for details, see Fig. S3). This correction step completed the segmentation and tracking of neurons in 3D space, and the information from the tracked neurons’ position was used to extract the activity of each neuron.

### 2.5 Results of extracting neuronal activities from three different test datasets

To evaluate the general applicability of our proposed method to 3D time-series images, we performed the segmentation and tracking using the 3D image series datasets from our original strain and from a previously published strain [19], both of which were obtained by using our own 3D imaging microscope system (dataset #1 and #2, respectively). We also analyzed a previously published 3D image series dataset [16] (dataset #3), which is different from dataset #1 and #2 in terms of strains, microscope system, and multiple aspects of image parameters (for details, compare panels A in Fig. 5, 6, S4, and S5). All worms were immobilized either by an anesthetizing drug or by constraining apparatus [16]; however, they still exhibited some movements that were solved using the tracking method. The neurons in these worms expressed red fluorescent protein (tdTomato [33], tagRFP [34], and mCherry [35] for datasets #1, 2, and 3, respectively) for a positional nucleus marker and genetically-encoded calcium indicator protein (GECI: GCaMP5G [36], GCaMP6s [37], and YC2.60 [38], respectively) in the nuclei for neuronal activity.

**Figure 5.**
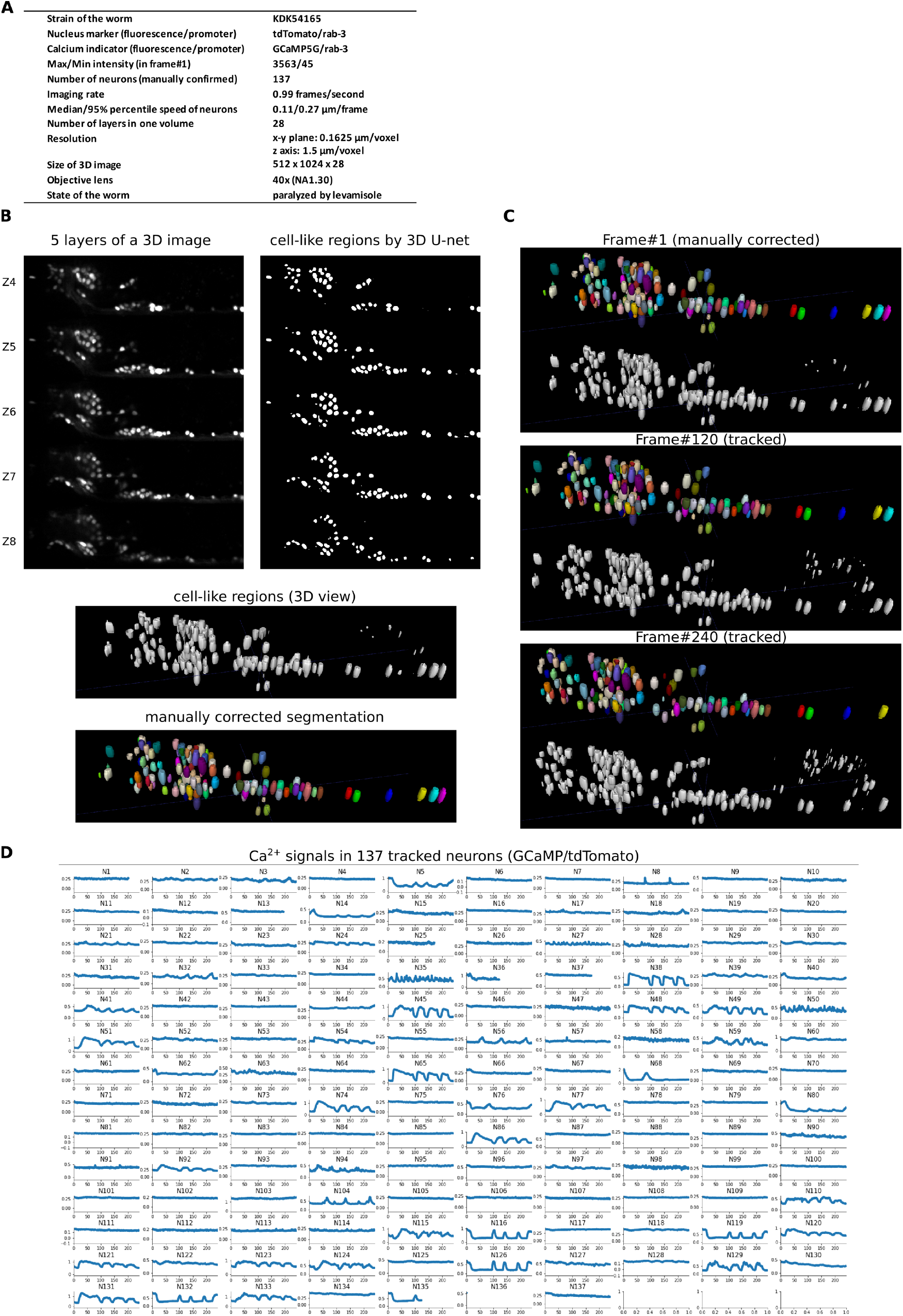
Results for an example type 1 dataset. A) Experimental conditions for this dataset. B) 3D image and its segmentation result in frame #1. Top left: Some of the layers of the 3D image. Top right: Cell-like regions corresponding to the figures at left, detected by the 3D U-net. Middle: Cell-like regions of the 3D view including all layers. Bottom: Final segmentations using watershed plus manual correction. C) Tracking results. Tracked cells in frame #120 and frame #240 are shown, which are transformations from the segmentation in frame #1. In each sub-figure, the top shows the tracked cells, and the bottom shows cell-like regions detected by 3D U-net. D) Extracted activities in 137 manually confirmed neurons. To eliminate movement artifacts, the activities are defined as the mean intensity of the GCaMP5G signal divided by the mean intensity of the tdTomato signal in each tracked cell region. The horizontal axis in each subfigure indicates the frame number, while the vertical axis indicates the activity. No neurons were mistracked; however, some neurons were lost because they moved out of the visual field of the camera.

We applied the above methods (Sections 2.1–2.4) on a test dataset of type 1 from the KDK54165 strain, which express tdTomato and GCaMP5G in most of the neurons. After segmentation, we manually confirmed 137 neurons in the head in the first frame of the 3D image series (Fig. 5B). We then applied the tracking method and successfully tracked all neurons, except for those that moved out of the field of view due to the worm’s motion (Fig. 5C, Movie S1). Interestingly, we found that many neurons showed dynamic spontaneous activities (Fig. 5D, Movie S2). We also found that many activities were synchronized with each other, e.g., N38 and N45 showed synchronized activities. This test dataset contained the fewest neurons of the type 1 datasets. We tested another type 1 dataset with more neurons (=164; KDK54165); however, that worm showed fewer neuronal responses (Fig. S4).

The same method was applied on a test dataset of type 2 from the AML14 strain [17]. Although this strain was very different than the KDK54165 strain in terms the nucleus marker intensity (268 vs. 3563) and inconsistent/vibrating movement (see Fig. S5A, 5A, and text below), our method still worked well (Fig. S5C, Movie S3)—We confirmed 101 neurons and successfully extracted activities for all neurons (Fig. S5D, Movie S4). It should be noted that fewer neurons were detected in the original report (about 90 or less) [17].

We applied the same method on a test dataset of type 3 using the JN2101 strain [16]. This dataset was obtained using a different strain, a different microscopy system, and under different experimental conditions than the type 1 and type 2 datasets (See Fig. 5A, 6A, S5A, and text below). As a consequence, this dataset had a lower resolution (half the resolution in the x and y directions) and larger displacements (about three times larger) between frames compared to datasets 1 and 2. However, our method worked well with a few modifications (see Method)—We confirmed and tracked 175 neurons in this dataset (marked by mCherry). Manual checking confirmed that only four neurons (N54, N62, N117, N122) had tracking errors (Fig. 6C, 6D, Movie S5), i.e. we correctly tracked 171/175 = 98% of the neurons. Our result was comparable to the original report in which 27 out of 198 neurons had tracking errors [16]. We extracted the neuronal activities from 19 neurons that expressed the GECI (Fig. 6D). We observed that neuron N108 showed a large response, and the response pattern was consistent with the previous report [16].

### 2.6 Challenging test conditions

To determine whether our method could be applied to even more difficult conditions, we tested our method using a series of generated datasets with increased difficulties by modifying the type 3 dataset. In all tests, we applied the 3D U-net as described before using a single training image from the original type 3 dataset. We also used the same manually corrected segmentation of the first frame based on the original image.

In the tracking task, one difficulty comes from large displacements of neurons between frames, and we increased this difficulty by removing intermediate frames in the image time-series. We generated and tested three datasets with 1/2 of the frames, 1/3 of the frames, and 1/5 of the frames as the original dataset. As expected, the number of tracking errors increased when more frames were removed (Fig. 7A, 7B). Even though the error rate in the 1/5 frame condition was rather high (25/175=14%, i.e. 86% correct), the error rate in the 1/3 frame condition was acceptable (14/175=8%), suggesting that our method could be applied to image time-series with much larger displacements than the type 3 dataset.

**Figure 6.**
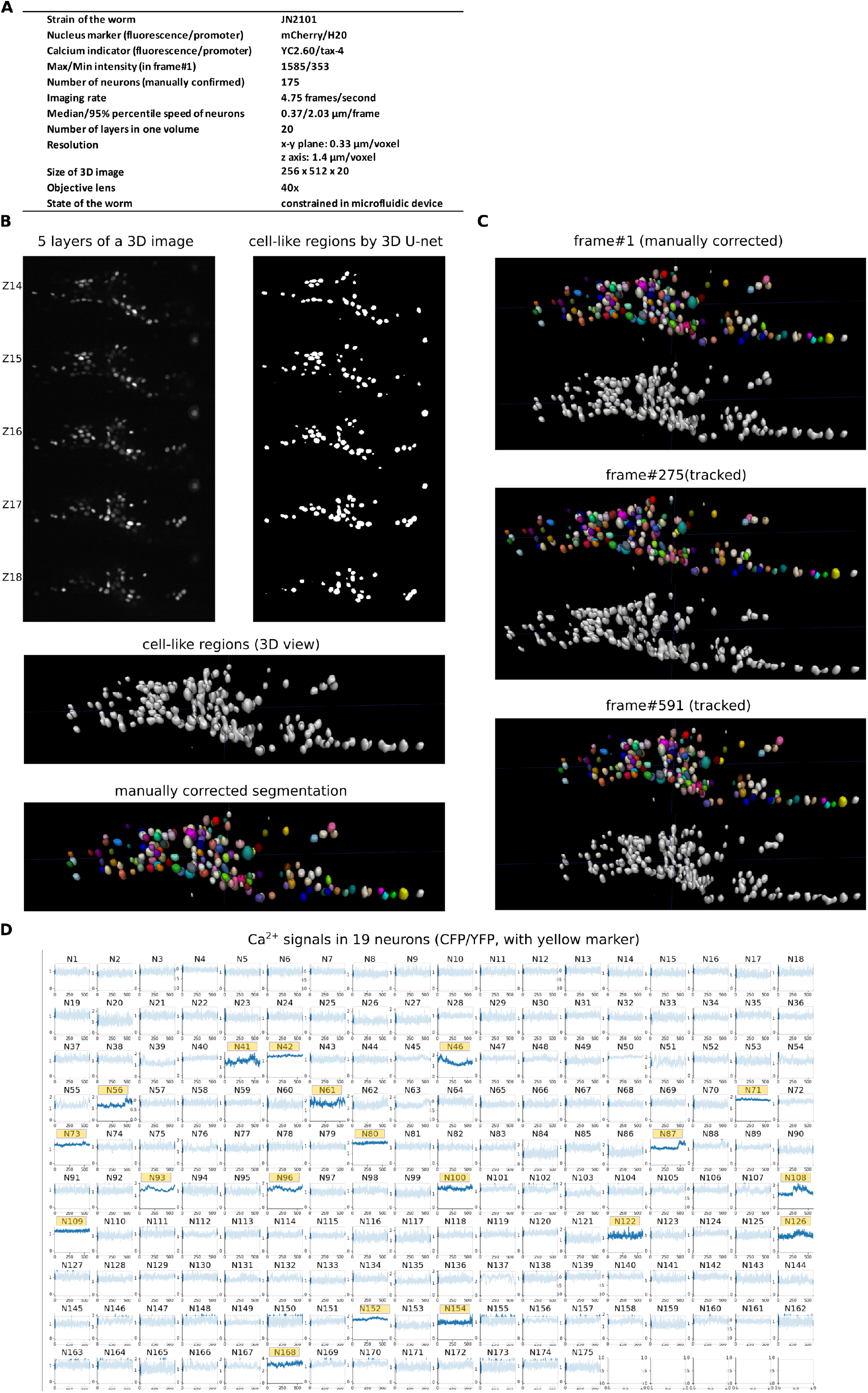
Results for a type 3 dataset. A) Experimental conditions for this dataset. B) 3D image and its segmentation result in frame #1. C) Tracking results. Tracked cells in frame #275 and frame #591 are shown. D) Extracted activities in 19 neurons (marked by yellow), expressing the calcium indicator YC2.60, which is driven by the tax-4 promoter. Signals in the other 156 neurons that do not express YC2.60 are artifacts. Only four neurons (N52, N62, N117, N122) were mistracked.

**Figure 7.**
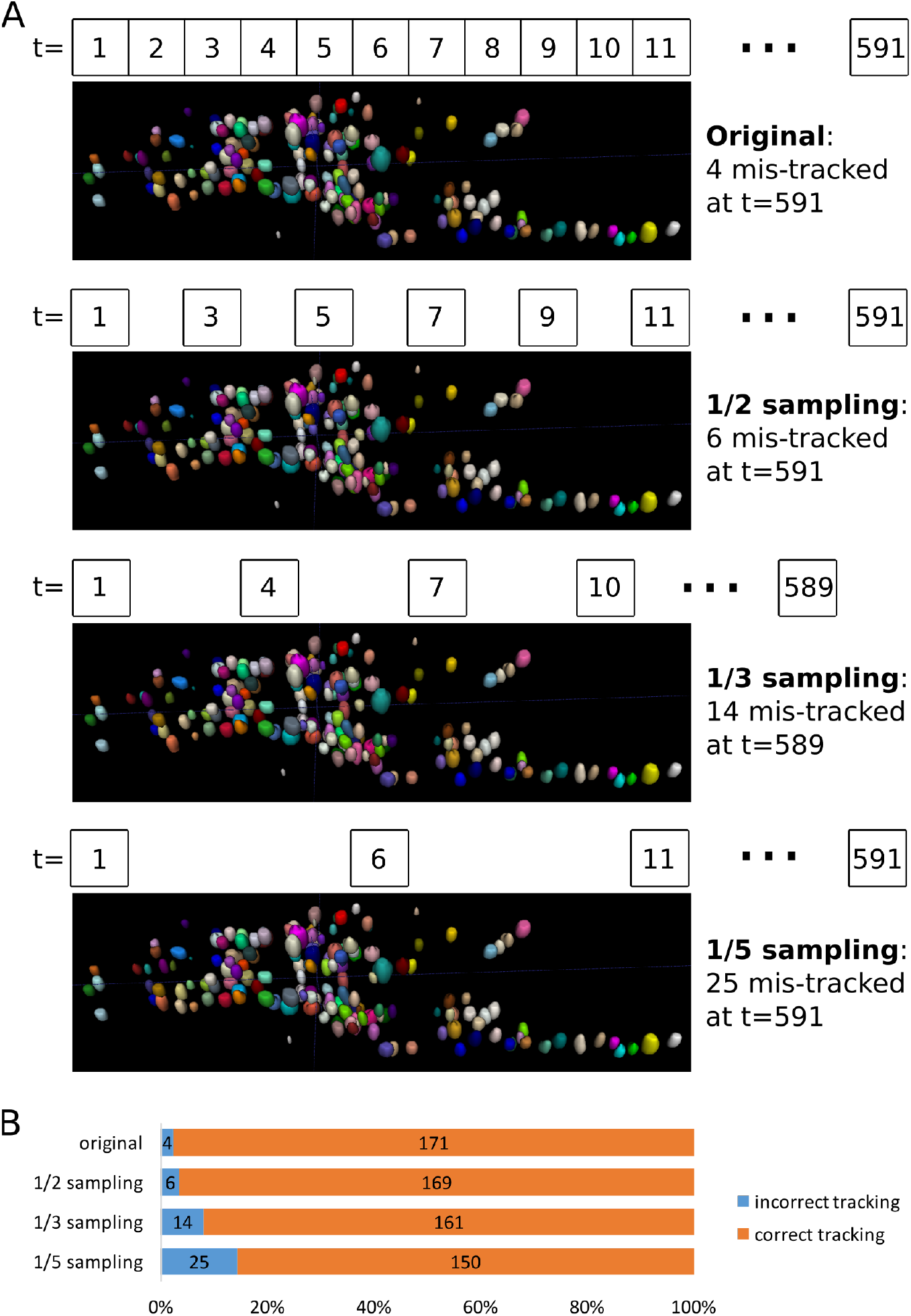
Testing the robustness of our method by removing intermediate frames from the type 3 dataset. A) Tracking results in the last frame at four different sampling rates. B) Bar graph showing the numbers of incorrectly tracked and correctly tracked neurons for the four sampling rates.

Another difficulty comes from the low signal-to-noise ratio in images. Large noise can obscure the real cell signal, thus leading to incorrect segmentation and tracking. We tested three modified datasets by adding different levels of Poisson noise into the original images. We added noise with levels of sd = 60, sd = 100, and sd = 140 (Fig. 8A, 8B), which was much larger than the noise level in the original images (sd = 4.05 in non-cell regions). In the sd = 60 condition, the image quality was obviously poorer than the original image, but our method still achieved a low error rate (6/175=3%). Even in the sd = 140 condition, where the image quality seemed to be very poor, our method achieved an acceptable error rate (16/175=9%, i.e. 91% correct). These results suggest that our method can be applied to datasets with very poor image quality when compared to the type 3 dataset. However, note that the segmentation in the first frame is manually corrected, and the manual process can also generate errors if the image quality is too low. Therefore, even though our tracking method was rather robust for low quality images, the image quality must be sufficient that we can correctly discriminate cells during the manual correction process.

**Figure 8.**
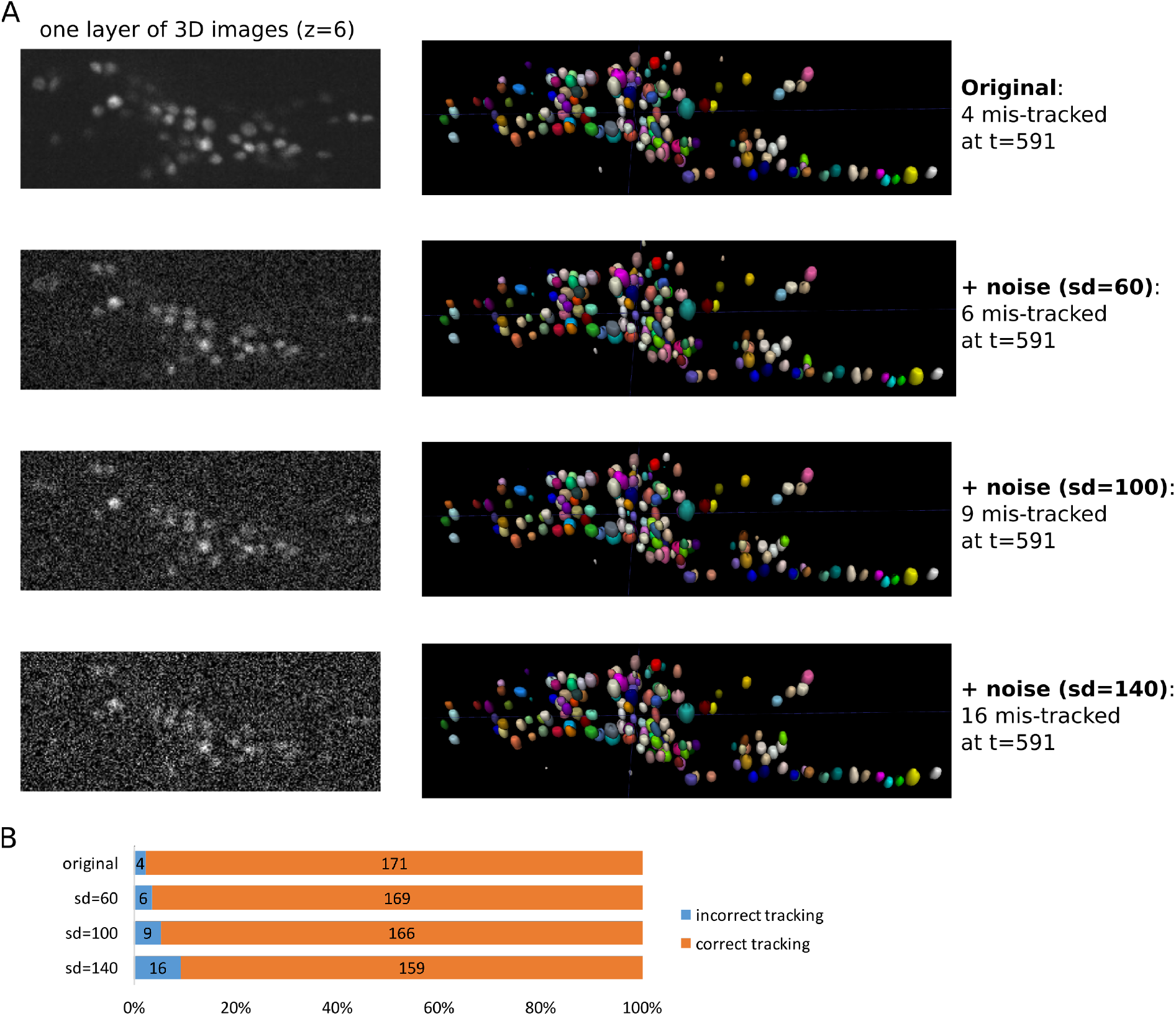
Testing the robustness of our method by adding Poisson noise into the type 3 dataset. A) Tracking results in the last frame for four different noise levels. B) Bar graph showing the numbers of incorrectly tracked and correctly tracked neurons for the four noise levels.

## 3. Discussion

We showed that our method was successfully applied to three different real datasets and to a series of more difficult generated datasets. We tested a variety of different conditions, including different nucleus marker fluorescence, cell level intensity, noise level, numbers of cells in each image, imaging rates, resolution or image sizes, and worm constraint conditions. Our method worked well in all of these different conditions with only slight modifications.

We also showed that our method can be easily modified to fit different conditions. When the resolution of the images was the same (datasets of type 1 and type 2), our method can be directly re-used without modification. When the resolution changed, we need to perform the following modifications: 1. Modify the structure of the 3D U-net; 2. Re-train the 3D U-net; and 3. Modify parameters in the program related to the resolution. Still, we only need to make a few modifications to the structure of the 3D U-net. For the re-training, the manual annotation usually take one day, and the network can be automatically trained in 1 or 2 h on our machine with a single GPU. Finally, several parameters related to the resolution, such as the minimum distance between cells and the minimum size of cells used in watershed, and the level of coherence in PR-GLS, should be modified accordingly (see Table 1). Thus, even when the imaging conditions are different and our method need to be adjusted, the number of parameters to be optimized are much smaller with our method than ones with the conventional methods (Table 1) due to the use of deep learning.

Because our tracking method tracked cells sequentially from the first frame to the subsequent frames, mistakes that occurred in one step will be maintained in all following frames. When the movement is small, the error rate is very low, so this is not problematic. However, with freely moving worms, the error rate was much higher, and a non-sequential strategy may be more robust. One possible approach to resolve this issue is to combine deep learning with ensemble methods, which construct a set of predictions (for segmentation and tracking) and vote based on all predictions. A previous study has applied a similar idea, but it did not use deep learning techniques [19]. One shortcoming of ensemble methods is that the correctness of segmentations and tracking cannot be guaranteed by the method itself. Therefore, the method will still need manual confirmation, which can be problematic when image quality is low, as we have described.

To further simplify the entire procedure, we may develop new network structures that combine more steps. This development may not be easy, but some recent studies have suggested possible directions. The U-net and 3D U-net are networks for semantic segmentation. Such networks classify each pixel as a specific category, such as human, animal, cell, or background. In contrast, there are networks designed for object segmentation (also called instance segmentation) that further separate objects in the same category into individuals: cell #1, cell #2, cell #3 and so on. Compared with semantic segmentation, object segmentation requires one more step and is thus more difficult to design the method, but this type of network would be suitable for our case. This field has also significantly advanced in recent years, and many methods have been developed [39–41]. These methods have been tested on public datasets such as common objects in context (COCO), but they have not been tested in cell segmentation. Furthermore, there is no current object segmentation method designed for 3D image segmentation. However, object segmentation may be one possible approach for improving cell segmentation in future studies. Another possible improvement could occur in the feedforward network—By designing a more appropriate structure and by using more training data, the feedforward network should give a better initial match that can be directly used for tracking all neurons.

In this study, we designed a new method to solve the segmentation and tracking problem for extracting whole-brain activities from 3D images. We used deep-learning techniques including the 3D U-net and a feedforward network, which can be flexibly modified for processing images taken under different conditions. By combining these deep-learning techniques with traditional image processing methods such as watershed and point set registration, we successfully segmented and tracked neurons from three different types of imaging datasets as well as from more difficult generated datasets. These results proved that our proposed method could be flexibly and widely applied to 3D whole-brain images obtained under different conditions.

## 4. Methods

### 4.1 Strains

The techniques used to culture and handle *C. elegans* were essentially the same as those described previously [42]. Both TQ1101 lite-1(xu7) and AML14 were obtained from the Caenorhabditis Genetics Center (University of Minnesota, USA). Young adult hermaphrodites were used in the imaging experiments.

### 4.2 Molecular biology and germline transformation

For pan-neuronal expression, NLS::tdTomato::NLS [43] and NLS::GCaMP5G::NLS (in which GCaMP5G [36] was codon-optimized for *C. elegans* and attached with NLS at N- and C-termini) were fused with *rab-3* promoter [44] using a GATEWAY system®(Thermo Fisher Scientific). Germline transformation into *lite-1(xu7)* [45] was performed using a microinjection [46] with a solution containing pYFU251 rab-3p::NLS::GCaMP5G::NLS (25 ng/μl), pYFU258 rab-3p::NLS::tdTomato::NLS (20 ng/μl), and OP50 genome (55 ng/μl) in order to obtain the strain KDK54165. The strain *lite-1(xu7)* was used to reduce the blue light-induced activation of the worm’s sensory system [45]. Independent transgenic lines obtained from the injection produced similar results.

### 4.3 Datasets

In this study, we used three dataset types. The 3D images in the type 1 and type 2 datasets were obtained using our custom-made microscope system, OSB3D (see Section 4.4). The type 1 worm strain was KDK54165, while the type 2 worm strain was AML14 [17,19] (wtfEx4[P*rab-3*::NLS::GCaMP6s: Prab-3::NLS::tagRFP]). The 3D images in the type 3 dataset were published in a previous report [16] and used worm strain JN2101 (Is[H20p::NLS4::mCherry]; Ex*tax-4*p::nls-YC2.60, *lin-44p*::GFP]).

### 4.4 OSB3D microscope system for the type 1 and type 2 datasets

We upgraded our robotic microscope system to 3D version [47]. For the 3D imaging, we used a custom-made microscope system that integrated the Nikon Eclipse Ti-U inverted microscope system with an LV Focusing Module and a FN1 Epi-fl attachment (Flovel, Japan). The excitation light was a 488-nm laser from OBIS 488-60 LS (Coherent) that was introduced into a confocal unit (CSU-X1) with a filter wheel controller (CSU-X1CU) to increase the rotation speed to 5,000 rpm (Yokogawa, Japan). The CSU-X1 was equipped with a dichroic mirror (Di01-T-405/488/561, Semrock) to reflect the 488-nm light to an objective lens (CFI S Fluor 40X Oil, Nikon, Japan), which passed through the GCaMP indicator used for calcium imaging and the red fluorescent protein used for cell positional markers. The laser power was set at 60 mW (100%). The fluorescence was introduced through the CSU-X1 into an image splitting optic (W-VIEW GEMINI, Hamamatsu, Japan) with a dichroic mirror (FF560- FDi01, Opto-line, Japan) and two bandpass filters (BA495-540 and BA570-625HQ, Olympus, Japan). The two fluorescent images were captured side-by-side on an sCMOS camera (ORCA Flash 4.0v3, Hamamatsu, Japan), which was controlled by a Precision T5810 (Dell) computer with 128-GB RAM using HC Image Live software (Hamamatsu) for Windows 10 Pro. A series of images for one experiment (about 1-4 min) required about 4-15 GB of space, which were stored on the 128-GB RAM during the experiment and transferred to a 1-TB USB 3.0 external solid state drive (TS1TESD400K, Transcend, Taiwan) for further processing.

For 3D imaging, the z-position of the objective lens was regulated by a piezo objective positioner (P-721) with a piezo controller (E665) and PIMikroMove software (PI, Germany). The timings of the piezo movement and the image capture were regulated by synchronized external edge triggers from an Arduino Uno (Arduino, Italy) using 35-ms intervals for each step, in which the image capture was 29.9 ms. For each step, the piezo moved 1.5 μm, and one cycle consisted of 29 steps. We discarded the top-most step because it frequently drifted off the correct position, and we used the remaining 28 steps. Note that one 3D image was 42 μm in length along the z-axis, which was determined based on the typical diameters of neuronal cell bodies (2-3 μm) and of a young adult worm’s body (30-40 μm). Each cycle took 1015 ms; thus, one 3D image was obtained per second. This condition was reasonable for monitoring neuronal activities because the worm’s neurons do not generate action potentials [48] and because many neuronal responses change on the order of seconds [49]. We also tested a condition using 10 ms for each step and 4.9 ms for an exposure using the same step size and step number per cycle (i.e., 2.3 frames of 3D images per second), which resulted in a comparable result (data not shown). For cyclic regulation of the piezo position, we used a sawtooth wave to assign positional information, instead of a triangle wave, because the sawtooth wave produced more accurate z positions with less variance among cycles.

### 4.5 Computational environment

Our image processing task was performed on a personal computer with an intel^®^ Core^TM^i7-6800K CPU @ 3.40 GHz × 12 processor, 15.5 GB of memory, and an Ubuntu 16.04 LTS 64-bit operating system. We trained and implemented the neural networks on a NVIDIA GeForce GTX 1080 GPU (8GB). The neural networks were constructed and implemented through the Keras high-level neural network API (https://keras.io), running on top of the Tensorflow machine-learning framework (Google, USA). All programs were implemented in python, except for the image alignment, which was implemented in ImageJ (NIH), and the image annotation and manual correction, which were implemented in ITK-SNAP (http://www.itksnap.org).

### 4.6 Pre-processing

Due to the movement of the worms, there could be small or large displacements between different layers of the same frame, which needed to be compensated for before the segmentation procedure. Using the StackReg plugin [50] in ImageJ (NIH), we compensated for the displacements by using rigid-body transformations to align each layer with the center layer. Neurons in the same worm can have very different intensities, which made it difficult to detect weak neurons. To solve this problem, we applied local contrast normalization [24] through a sliding window (27 × 27 × 3 pixels) so that all neurons had similar intensities. This normalization was applied only to the nucleus marker images and did not affect to calculate the signal intensities for calcium imaging.

### 4.7 3D U-net

We used a 3D U-net structure similar to that shown in the original study [25]. The network received a 3D image as input and generated a 3D image of the same size with values between 0 and 1, indicating the probability that each pixel belonged to a cell region. The 3D U-net structure shown in Fig. 3A was used on our type 1 and type 2 datasets, which have the same resolution. For the type 3 dataset, which had a lower resolution and smaller size, we reduced the number of pooling operations and the sizes of the input, output, and intermediate layers. The smaller structure occupied less GPU memory, so we increased the number of convolutional filters on each layer to increase the capacity of the network (see Fig. S1).

The U-net can be trained using very few annotated images [28]. In this study, we trained two 3D U-nets: one for the type 1 and 2 datasets and one for the type 3 dataset. Each 3D U-net used one 3D image for training. The image was manually annotated into cell regions and background regions using the ITK-SNAP software. We used the binary cross-entropy as the loss function to train the 3D U-net. Because the raw image sizes were too large (512 × 1024 × 28 or 256 × 512 ×20), we divided the raw images into small sub-images that fit the input sizes of the two 3D U-nets (160 × 160 × 16 or 96 × 96 × 8). To improve the 3D U-net performance, we used dataset augmentation to increase the training data by applying random affine transformations to the annotated 3D images. The affine transformation was restricted on the x-y plane but not in z-direction because the resolution in the z-direction is much lower than that in the x-y plane (see Fig. 5A, 6A, and S5A). To verify the generalization of our 3D U-net, we used test datasets that were independent of the type1 and type 2 training datasets. Because we had only one type 3 dataset, we trained the 3D U-net using the frame #1 image and applied the 3D U-net to all frames of the dataset.

### 4.8 Watershed

The 3D U-net generated probability outputs between 0 and 1 that indicated the probability a pixel belonged to a cell-like region. By setting the threshold to 0.5, we could divide the 3D image into cell-like regions (>0.5) and background regions (≤0.5). The cell-like regions in the binary images were further transformed into distance maps, where each value indicated the distance from the current pixel to the nearest background region pixel. We applied a Gaussian blur to the distance map and searched for local peaks, which were assumed to be cell centers. We then applied watershed segmentation [26] using these centers as seeds. Watershed segmentation was applied twice; the first application was 2D watershed segmentation in each x-y plane, and the second application was 3D watershed segmentation of the whole image. Two segmentations were used because the resolution in the x-y plane and z-direction differed, and we needed to establish different minimal distances between local peaks.

### 4.9 Feedforward network

As Fig. 4B illustrates, the input of the network contained the position information for two points. Each point was represented by the normalized positions of the 20 nearest neighboring points. The 20 nearest neighbor positions 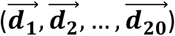 were given by 3D vectors because they were extracted from the 3D image. To normalize the points, each of the 20 positions was divided by the mean distance 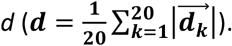 The normalized positions were then sorted by their absolute values, from shortest to longest. Finally, the mean distance *d* was included as the last value, so each point was represented by a 61D vector.

We used the first fully connected layer to calculate the learned representation of the relative positions of each point as a 512D vector (Fig. 4B, the first hidden layer after input). The vectors for the two input points were concatenated to become a 1024D vector. We then applied a second fully connected layer on this 1024D vector to compare the representations of the two points. The resulting 512D vectors were processed by a third fully connected layer to obtain a single similarity score between 0 and 1, which indicated how likely that two points came from the same neuron.

We performed segmentation on a single frame of the type 3 dataset and obtained a point set for the centers of all neurons. Because we needed a large number of matched point sets for training and manually matching point sets is time-consuming and impractical, we created an artificial training dataset by applying random affine transformations to the point set described above and adding small random movements to each point. The correspondence between the generated point sets was completely known, so we could train the feedforward network with the artificially generated dataset.

We used binary cross-entropy as the loss function to train the feedforward network.

### 4.10 PR-GLS method

The PR-GLS method requires an initial match to start the tracking process [31]. This initial match was calculated using the similarity scores from the feedforward network. In this procedure, we matched the two points from two different frames with the highest scores, deleted these two points, and matched the next set of two points with the highest scores. By repeating this process, we obtained an initial match (Fig. 4A). This initial match was corrected using the expectation-maximization (EM) algorithm in the PR-GLS method as described in the original paper [31]. We set the maximum number of EM iteration steps to be 20 and obtained satisfactory results in all test datasets. In the original paper [31], the initial match (by FPFH) was re-calculated during the EM iterations, but we only calculated the initial match once (by feedforward network) before the EM steps were performed without problems. After the PR-GLS corrections, we obtained coherent transformations from the points of each frame to the subsequent frame (Fig. 4A).

### 4.11 Extracting activities

After we tracked the manually confirmed neurons from the first frame to the last frame, we could easily extract activities from the regions corresponding to each neuron. We measured the intensities in both channels corresponding to the Ca^2+^ indictor and the nucleus marker. The activity was computed as GCaMP5G/tdTomato in the type 1 dataset, GCaMP6s/tagRFP in the type 2 dataset, and CFP/YFP in the type 3 dataset.

### 4.12 Manual check and correction

We manually checked the results of segmentation and tracking. For segmentation, we superimposed the automatically segmented regions of frame #1 onto the raw 3D image using the ITK-SNAP software. By careful observation, we discarded artifact regions, such as autofluorescence regions and neuronal processes. Over-segmentation and under-segmentation regions were corrected according to the size and shape of normal neurons. For cell tracking, we combined two 4D images, namely the tracked labels and the raw image time series, in a top-bottom arrangement in the ImageJ software in order to compare the images at each time point. By observing the correspondence between the tracked labels and the raw images at different time points, we could identify mistakes in the tracking result.

## Acknowledgement

We are deeply grateful to those people for their advices, comments and technical assistances for this study, including Toru Tamaki, Ichiro Takeuchi, Katsuyoshi Matsushita, Hiroyuki Kaneko, Taro Sakurai, and members of the Kimura lab. Nematode strains were provided by the Caenorhabditis Genetics Center (funded by the NIH Office of Research Infrastructure Programs P40 OD010440). Funding:

This study was supported by a grant by the Ministry of Education, Science and Technology (KAKENHI JP16H06545) (KD.K.), a program for Leading Graduate Schools entitled ‘Interdisciplinary graduate school program for systematic understanding of health and disease’ (T.M.), and RIKEN Center for Advanced Intelligence Project.

---

**Figure S1.**
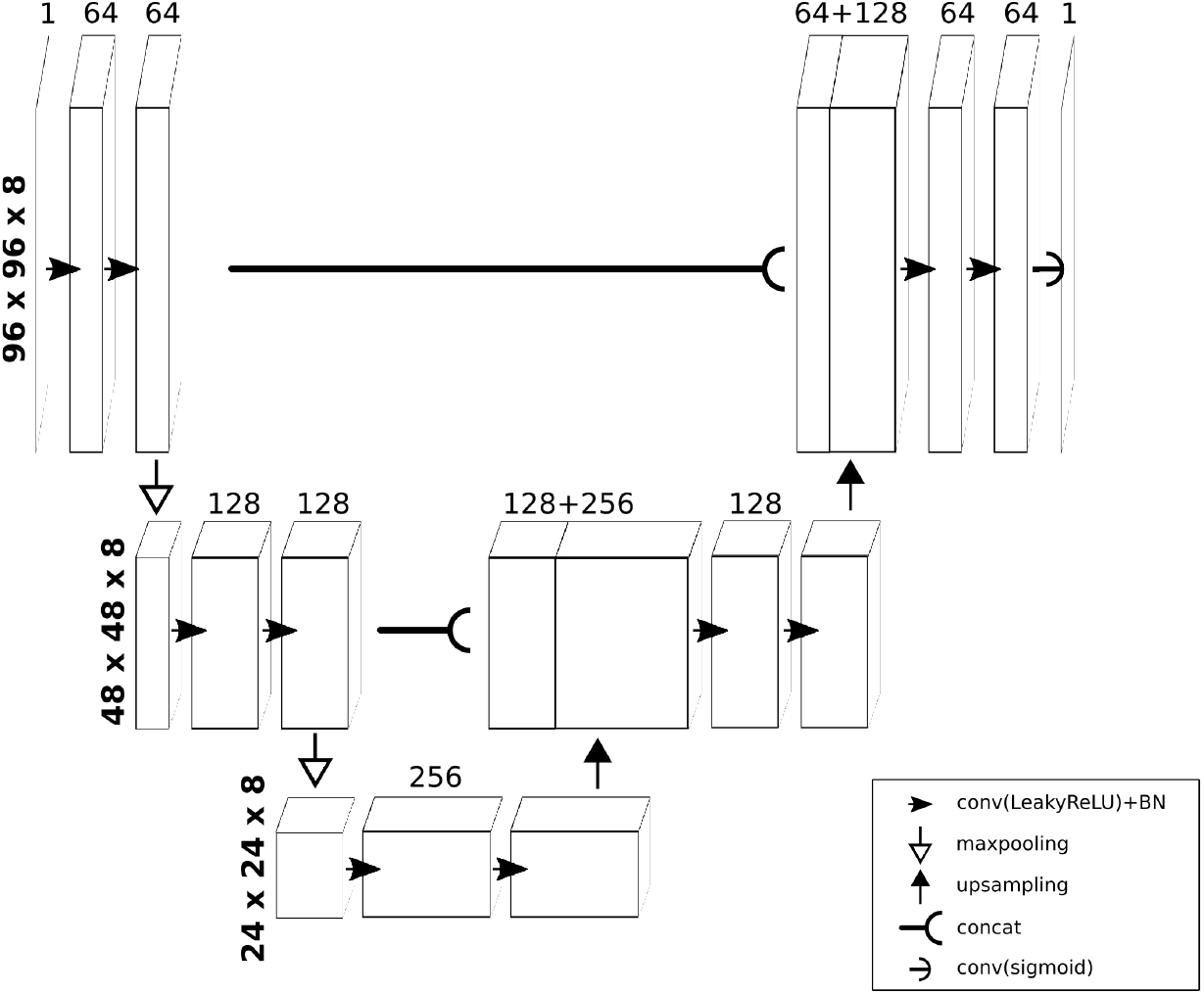
Structure of the 3D U-net used for the type 3 dataset.

**Figure S2.**
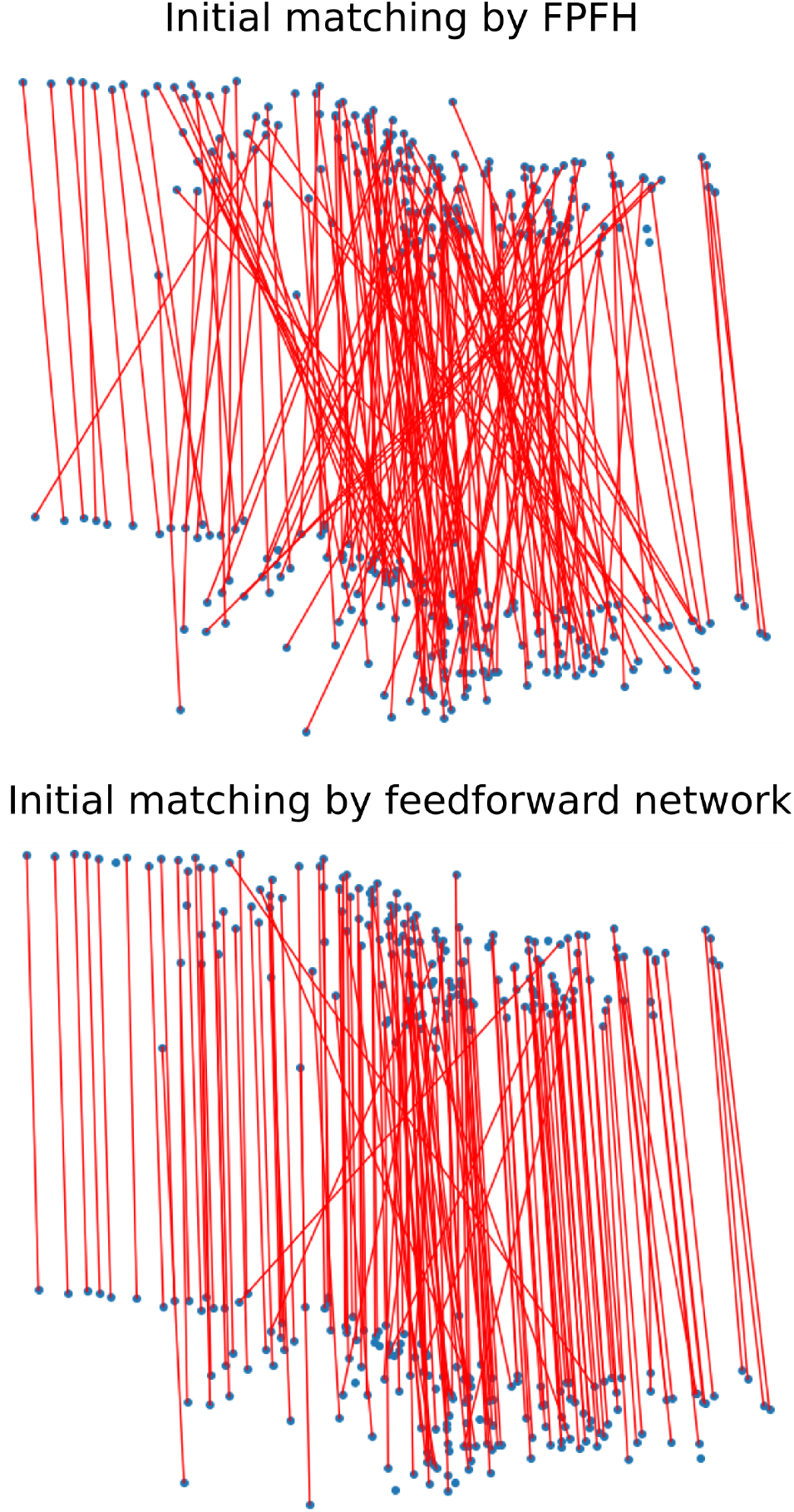
Examples of initial matchings by FPFH and our feedforward network. In each subfigure, there are two point sets (blue points at top and bottom) from two different frames of the same worm. Red lines indicate the estimated matchings between these two sets.

**Figure S3.**
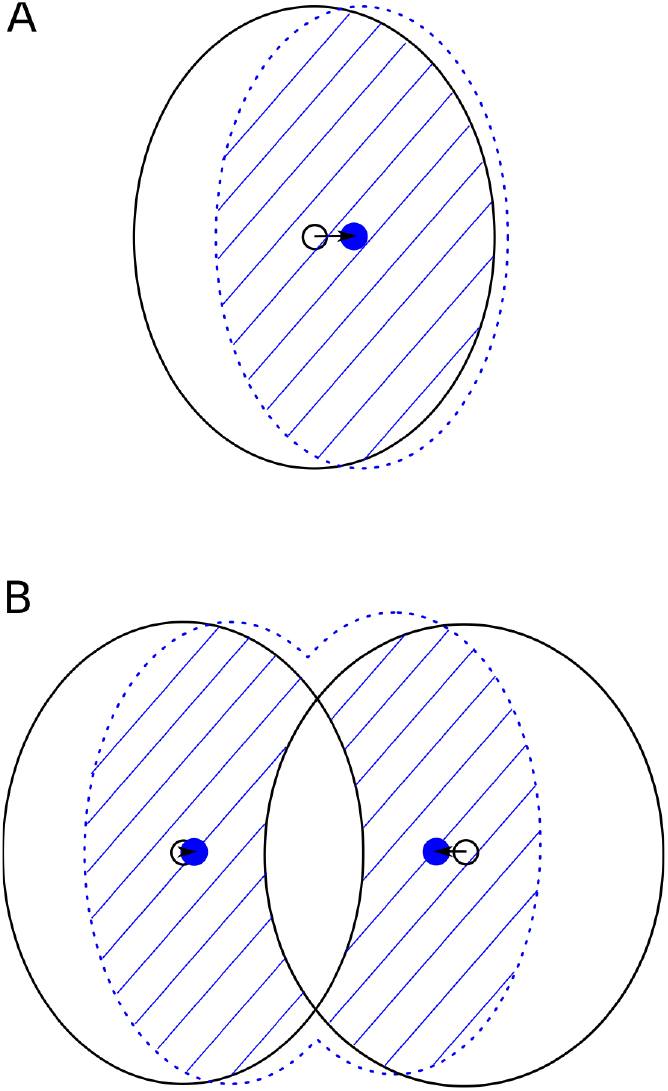
Accurate correction of the coherent transformation. A) Correction method in the single neuron case. The large black ellipse is the region of a cell predicted by PR-GLS, while the small black circle is the center of the PR-GLS predicted region. The dashed blue ellipse is the region detected by 3D U-net. The region filled with blue lines indicates the intersection of the PR-GLS region and the 3D U-net region, and the small blue circle indicates the center of the intersection region. The arrow shows the correction vector suggested by this method, i.e., the center of the cell should move from the small black circle to the small blue circle. B) Correction method in the case that two predicted regions partially overlap. The method is basically the same as in A), except the overlapped region between two black ellipses is discarded when the intersection regions (blue lines) are determined in order to prevent merging of the two neurons.

**Figure S4.**
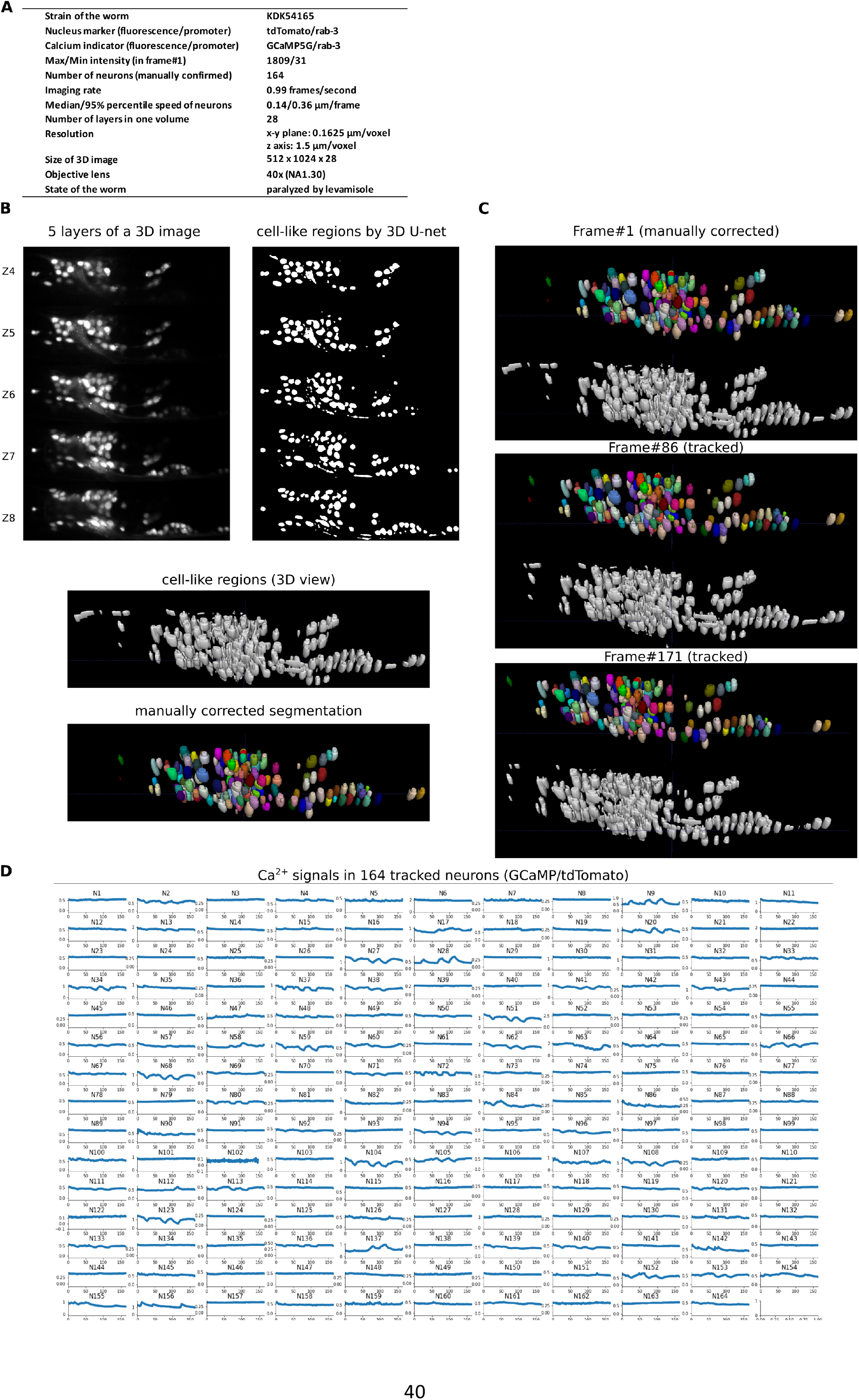
Results for another type 1 dataset. A) Experimental conditions of the dataset. B) 3D image and its segmentation result in frame #1. C) Tracking results. Tracked cells in frame #86 and frame #171 are shown. D) Extracted activities in 164 manually confirmed neurons. No neuron was incorrectly tracked.

**Figure S5.**
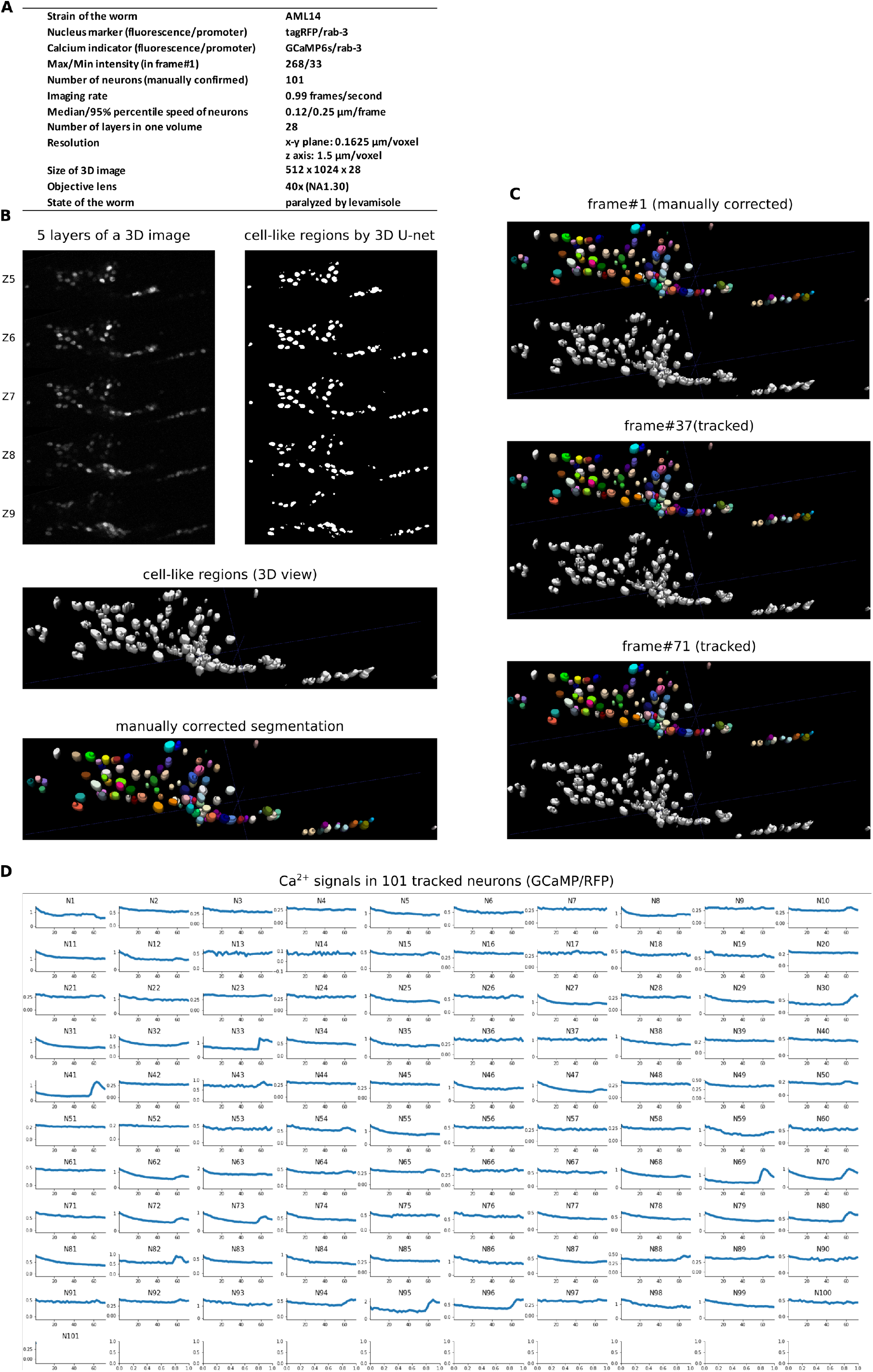
Results for the type 2 dataset. A) Experimental conditions of the dataset. B) 3D image and its segmentation result in frame #1. C) Tracking results. Tracked cells in frame #37 and frame #71 are shown. D) Extracted activities in 101 manually confirmed neurons. No neuron was incorrectly tracked.

**Table 1.** Comparison of the complexities between the previous method [16] and our method. For each method, the procedures are listed along with the number of parameters (red) utilized.

|  | Toyoshima *et al.* 2016 |  | Our method |  |
| --- | --- | --- | --- | --- |
|  | Procedures | Parameters | Procedures | Parameters |
| Pre-processing | Alignment | - | Alignment | - |
|  | Denoising (median filter) | (1) Radius | Local contrast normalization<br>(make all cells have similar intensities) | (1) Noise level |
|  | Background subtraction | (2) Radius |  |  |
|  | Gaussian blurring | (3) Radius |  |  |
| Segmentation | Thresholding | (4) Method (e.g. 'mean', 'triangle') | 3D U-net<br>(detect cell/non-cell regions) | Automatically learned |
|  | Watershed | (5) Radius<br>(6) Minimum size of cells |  |  |
|  | Counting number of negative curvature | (7) Threshold<br>(8) Distance to borders | Watershed (twice) | (2) Minimum distance,<br>(3) Minimum size of cells |
|  | Watershed 2 | - |  |  |
| Tracking | Least squares fitting of Gaussian mixture | (9-11) Default value of covariance matrix (3d vector) | Feedforward network<br>(generate an initial matching) | Automatically learned |
| | | | PR-GLS (generate a coherent transformation from the initial matching) | (4, 5) coherence level controlled by $\beta$ and $\lambda$<br>(6) maximum iteration |
|  | Removing over-segmentation | (12) Distance | Accurate correction using intensity information | - |

